# SNRK regulates TGFβ levels in atria to control cardiac fibrosis

**DOI:** 10.1101/2024.09.24.612951

**Authors:** Karthikeyan Thirugnanam, Farhan Rizvi, Arshad Jahangir, Peter Homar, Fathima Shabnam, Sean P. Palecek, Suresh N. Kumar, Amy Pan, Xiaowen Bai, Hidekazu Sekine, Ramani Ramchandran

## Abstract

Atrial fibrosis is central to the pathology of heart failure (HF) and atrial fibrillation (AF). Identifying precise mechanisms underlying atrial fibrosis will provide effective strategies for clinical intervention. This study investigates a metabolic serine threonine kinase gene, *sucrose non-fermenting related kinase (SNRK),* that we previously reported to control cardiac metabolism and function. Conditional knockout of *Snrk* in mouse cardiomyocytes (*Snrk* cmcKO) leads to atrial fibrosis and subsequently HF. The precise mechanism underlying cardiomyocyte SNRK-driven repression of fibrosis is not known. Here, using mouse, rat, and human tissues, we demonstrate that SNRK expression is high in atria, especially in atrial cardiomyocytes. SNRK expression correlates with lower levels of pro-fibrotic protein transforming growth factor-beta 1 (TGFβ1) in the atrial cardiomyocytes. In HL-1 adult immortalized mouse atrial cells, using siRNA approaches, we show that *Snrk* knockdown cells show more TGFβ1 secretion, which was also observed in heart lysates from *Snrk* cardiac-specific knockout mice in vivo. These effects were exacerbated upon infusion of Angiotensin II. Results from *Snrk* knockdown cardiomyocytes co-cultured with cardiac fibroblasts suggest that SNRK represses TGFβ1 signaling (Smad 2/3) in atrial CMs and prevents paracrine cardiac fibroblast activation (α-SMA marker). In conclusion, high SNRK expression in atria regulates cardiac homeostasis, by preventing the release of TGFβ1 secretion to block cardiac fibrosis. These studies will assist in developing heart chamber-specific fibrosis therapy for non-ischemic HF and AF.

## Introduction

Heart failure (HF) is the final consequence of an adverse cardiovascular event. Although progress has been made in treating HF, hospitalization and prevalence rates continue to rise, thus requiring more research into the causes of HF and associated arrhythmias, and ways to prevent and treat HF (Borlaug and Paulus, 2011; Dick and Epelman, 2016; Giamouzis et al., 2010; Ponikowski et al., 2014). HF is associated with cardiac remodeling, a post injury cardiac repair process in which inflammation and fibrosis processes play key roles. Both these processes are reversible in pre-clinical and clinical settings, which forms the underlying basis for HF treatment options (Díez et al., 2002; Brilla et al., 2000). However, the pathogenesis of inflammation and fibrosis associated with ischemic vs. non-ischemic HF are distinct (Bacmeister et al., 2019). For example, in ischemic HF, inflammation precedes fibrosis whereas in non-ischemic HF, inflammation and fibrosis co-occur. Both inflammation and fibrosis are also important processes for normal cardiac function and pathogenesis of cardiac arrhythmias (Mehta, and Griendling, 2007; Wu et al., 2017). Thus, strategies to mitigate HF that can selectively interfere with one process but not the other will be beneficial. Data from our team shows that inflammation and fibrosis during non-ischemic HF can be decoupled (Thirugnanam et al., 2019), and thus provides a novel avenue for the development of selective intervention strategies for non-ischemic HF. Fibrosis in the atria and ventricles of the heart can have different effects on cardiac function. Atrial fibrosis is often associated with or leads to atrial fibrillation (AF), a clinical condition more common during cardiac arrhythmia (Sygitowicz et al, 2021). The prevalence of AF in patients with HF increased with the severity of the disease (Maisel, and Stevenson, 2003). Atrial AMP-activated protein kinase (AMPK) is critical for prevention of dysfunctional electrical excitation and AF (Su et al., 2022). Therefore, if targets in the AMPK family exist that show preferential expression in the atrium vs. ventricular chamber, then strategies targeting such signaling will offer some selectivity, and compounds generated with this approach are likely to have minimal side effects (Rizvi et al., 2016). Thus, the need for identifying such targets and the underlying mechanisms associated with preventing or promoting fibrosis.

Our team and others have been studying the role of sucrose non-fermenting related kinase (SNRK), an AMPK family member in the cardiac system (Thirugnanam et al., 2019; Cossette et al., 2016; Cossette et al., 2014; Thirugnanam and Ramchandran, 2020; Rines et al., 2017) for several years, that may serve as one such target for selective modulation of cardiac fibrosis. Recently, a circular RNA for *SNRK* was identified, that targets microRNA (*miR-103-3p*) to promote cardiac repair in rats with myocardial infarction (Zhu et al., 2021). *Snrk* is essential for cardiac metabolism and function, effects that are cardiomyocyte (CM) cell autonomous (Cossette et al., 2016; Cossette et al., 2014). Using a chronic infusion Angiotensin II (Ang II) mouse model of non-ischemic HF, we have shown that CM-specific *Snrk* knockout mice (*Snrk* cmcKO) are susceptible to HF within 2 weeks post-infusion (Thirugnanam et al., 2019) compared to their control littermates. Interestingly, endothelial-specific *Snrk* knockout mice (ecKO) in the same Ang II model are not susceptible to HF compared to control littermates. Further, under basal state, *Snrk* cmcKO hearts show enhanced fibrosis while *Snrk* ecKO hearts do not, and upon Ang II infusion, enhanced fibrosis is observed in both conditions (Thirugnanam et al., 2019). Whether fibrosis induction in the atria and ventricle of the heart are similar in *Snrk* cmcKO mice is not known. We sought to investigate this question in this study. Previous work showed that overexpression of a mutant (C33S) form of constitutively active transforming growth factor-β1 (TGF-β1) expression under cardiac restricted α-myosin heavy chain promoter showed selective atrial chamber specific fibrosis but not ventricular chamber fibrosis even though the protein was expressed constitutively in both chambers (Nakajima et al., 2000). The underlying mechanism for this selective atrial fibrosis in this mutant TGF-β transgenic overexpressing mice was not known (Nakajima et al., 2000). Our work here sheds light on SNRK’s role in mediating chamber-specific cardiac fibrosis and the underlying mechanism associated with it.

## Methods

### Animal work

#### Mice

The mice were housed in the Medical College of Wisconsin Biological Resource Center, and all experiments were performed following an approved IACUC animal protocol 1022. The *Snrk cmcKO* (cardiac myocyte knockout) adult mice were generated from *MYH6CRE-positive Snrk LoxP/wild-type (WT)* males mated to *Snrk LoxP/LoxP* females. The genotyping and characterization information for these alleles are available from our previous publication (Cossette et al., 2014). Please note genotyping data in Fig. S7 in Cossette et al, Biology Open publication and SNRK siRNA efficacy in HL-1 cells in Fig. S2 in Thirugnanam et al 2019 JAHA publications and Fig. 4 in this manuscript. The in vivo experiments included the following conditions: *Snrk WT, Snrk cmcKO, Snrk WT* with Ang II, and *Snrk cmcKO* with Ang II. Ang II osmotic pump infusion studies at a dose of 1000 ng/kg/minute were performed as described before (Thirugnanam et al., 2019). Additional surgical details and recovery are included here. The experimental animals for the data point were all males in the study and n=3 for each condition. We initially began the experiment with 5-7 mice per group but due to mortality associated with the Ang II infusion, we had only 3 mice remaining at day 14 for analysis.

#### Rat

All animal experiments were performed following the guidelines of the Ethical Committee for Animal Experimentation of Tokyo Women’s Medical University and in compliance with the laws and regulations on the use of animals in biological research and the ARRIVE guidelines. All experimental protocols were approved by the Animal Experimentation Committee of Tokyo Women’s Medical University as AE23-104. All animals were housed in individual cages with free access to food and water under a 12-h light/dark cycle and maintained at constant room temperature and humidity. Animals were euthanized by exsanguination under 5% isoflurane following the American Veterinary Medical Association (AVMA) euthanasia guidelines. Sprague-Dawley rats (aged 8 weeks; male) were obtained from Japan SLC, Inc., Japan.

#### Ang II infusion studies (rat)

This study was carried out using hearts from 12-week-old rats weighing 300-350 g, purchased at 8 weeks and weighing 300-320 g. Rats were anesthetized and subcutaneously implanted with an osmotic mini pump (ALZET Osmotic Pumps 2ML4, Cupertino, CA, USA) that released Angiotensin II (KareBay Biochem, Monmouth Junction, NJ, USA) at a constant rate of approximately 1.4 mg/kg per day for 4 weeks to create a model of cardiac inflammation and fibrosis.

#### Ang II infusion studies (mice)

To simulate cardiac stress, osmotic pumps that secrete a steady flow of angiotensin II were used. The pumps were implanted at 4 months of age and transthoracic ECHO analysis were performed as described before (JAHA 2019) immediately prior to implantation followed by analysis once per week for a total of 2 to 4 weeks. At the end of the experiment the mice were euthanized using CO_2_ and cervical dislocation.

#### Osmotic pump install surgery

For this procedure, anesthesia was induced by placing the mice into an induction chamber using 3-5% isoflurane. Once the mice were anesthetized, they were transferred to the heated platform, and anesthesia with isoflurane was maintained at a concentration of 1.5-2% using a nose cone. During anesthesia, the mid-scapular region of the anesthetized animal was shaved.

#### Osmotic pump implantation

Either betadine or chlorhexidine scrub was used followed by an alcohol prep of the site to disinfect the skin prior to incision. A small incision was made in the skin. A sterile hemostat was inserted into the incision and gently opened and closed to create a subcutaneous pocket for the pump. The drug-filled pump (freshly cleaned with alcohol wipe) was inserted (delivery port first) into the pocket. The incision was closed with wound clips. The animal was placed on soft paper towels to allow recovery from anesthesia. The animal was monitored continuously following surgery until they were awake and able to walk around normally. We inject subcutaneously once Buprenorphine-SR as an analgesic due to its long-acting effects. Effects of one injection last for 72 hrs. Analgesic is given under anesthesia prior to recovery. Wound clips were removed 10-14 days after pump placement.

#### Human studies

The study was approved by the Aurora Institutional Review Board and conforms to the Health Insurance Portability and Accountability Act of 1996. All the non-heart failure samples were obtained from the National Disease Research Interchange. Philadelphia PA. All the heart failure samples were obtained from heart transplant patients and all of them were with reduced ejection fraction (EF<45). Atrial and ventricular tissue were dissected and lysed in RIPA buffer for western blotting analysis. Details of the patient information are available in supplemental figure S3G.

### Immunohistochemistry

#### Macrophage staining

Formalin-fixed hearts collected from *Snrk WT* and *Snrk cmcKO* mice treated independently with vehicle and Ang II were prepared as described previously (Thirugnanam et al., 2019). Briefly, for assessing macrophage infiltration in hearts, anti-mouse/human Mac2 (Galectin-3) antibody (Cedarlane, Burlington, ON, Canada; catalog No. CL8942AP) was used. Pretreatment with target retrieval solution (Dako, Santa Clara, CA; S1699) was performed with heating (95°C) over a period of 40 min and then cooling at room temperature for 15 min. Tissue slices were incubated overnight at 4°C, and stained sections were viewed using light microscopy. All analyses of histological data quantification were done by scanned images that were imported to Visiopharm software (Hørsholm, Denmark) and using the imager module; 10 ×20 region-of-interest images were extracted from left, center, and right regions of the heart. All original images were processed with this preset threshold and linear Bayesian classification to generate the processed image.

#### Fibrosis studies

*Picros*irius red staining was performed to determine the extent of fibrosis in the heart sections. The amount of fibrosis was quantified using a software-assisted, unbiased microimage quantification method. Image analysis was fully blinded without the analyst’s knowledge of the groups and treatments. The scanned images were imported into Visiopharm software (Hørsholm, Denmark), and the region of interest was marked in each of the heart’s left, center, and right ventricular walls. Similarly, the left and right atrium wall areas were also marked for analysis. All original images were processed at 20x magnification with a preset threshold to estimate the fibrotic region (red stain), and linear Bayesian classification was performed to generate the processed image. The perivascular fibrotic region was identified using the lumen in the center and eliminated by a post-processing filter. The total interstitial collagen-positive area per region of interest was measured in micrometers and represented as a percentage of the total tissue area.

#### Alpha smooth muscle actin staining

Briefly, formalin-fixed hearts collected from WT and *Snrk* cmcKO mice treated independently with vehicle and Ang II were dehydrated, embedded in paraffin, and sectioned at 5 μm. Slides were warmed at 60°C for 1 h, dewaxed in xylene, and rehydrated. For assessing alpha smooth muscle actin expression, anti-alpha smooth muscle actin antibody (Abcam, Cat# 5694, 1:200 dilution) was used. Pretreatment with target retrieval solution (Dako, Santa Clara, CA; S1699) was performed with heating (95°C) over a period of 40 min and then cooling at room temperature for 15 min. Tissue slices were incubated overnight at 4°C, and stained sections were viewed using light microscopy. All analyses of histological data quantification were done by scanned images that were imported to Visiopharm software (Hørsholm, Denmark) and using the imager module; 10 ×20 region-of-interest images were extracted from left, center, and right regions of the heart. All original images were processed with this preset threshold and linear Bayesian classification to generate the processed image.

#### Immunofluorescence studies

For cross-sectional observation, rat hearts were fixed in 4% paraformaldehyde and processed into 7 µm-thick paraffin-embedded sections. For detection of SNRK, deparaffinized sections were incubated with a 1:100 dilution of anti-SNRK rabbit polyclonal antibody (Genetex, Irvine, CA, USA). For detection of cardiomyocytes, sections were incubated with 1:100 dilution of anti-cardiac troponin T rabbit polyclonal antibody (Abcam, Cambridge, UK) and for detection of TGFβ protein, sections were incubated with 1:100 dilution of anti-TGFβ rabbit polyclonal antibody (Genetex) at 4°C overnight at incubated overnight at 4°C. The sections were then incubated with a 1:200 dilution of Opal Polymer anti-Rabbit HRP (Akoya Biosciences, Marlborough, MA, USA) for 1 h at room temperature. Both the primary and secondary antibodies were diluted in 1X PBS. Signals were generated using Opal Fluorophore (Opal520, 570, Akoya Biosciences) and nuclei were counterstained with Spectral DAPI (Akoya Biosciences) for 5 min at room temperature. Sections were finally visualized using a confocal laser scanning microscope (FV1200: Olympus, Tokyo, Japan).

#### Cell lines, transfection, and co-culture studies

Mouse HL-1 atrial cardiomyocytes were purchased from Sigma Aldrich (Cat# SCC065). HL-1 cell line was cultured in Claycomb media (Sigma-Aldrich, Cat# 51800C) containing 10% FBS, 0.1 mmol/L norepinephrine (Sigma-Aldrich, Cat# A0937), 2 mmol/L l-glutamine (Sigma, Cat# G7513, and penicillin/streptomycin (Thermo Fisher, Cat# 15070063). The culture plates were precoated with gelatin/fibronectin overnight at 37°C before seeding the cells. Mouse cardiac fibroblasts were purchased from Sciencell (Cat# M6300-57) and cultured in Fibroblast medium-2 (Sciencell, Cat# 2331). The culture plates for fibroblasts were precoated with Poly-L-Lysine (Sciencell, Cat# 0413) for 1 h prior to seeding the cells. Knockdown in HL-1 cells was performed with small interfering RNA (siRNA) for *Control* scrambled (25 μM) (Cat# D-001320-10-05) and *Snrk* (Cat# J-051065-05-0005) (25 μM) from Horizon-inspired cell solutions. Briefly, HL-1 cells were seeded in 6-well plates approximately 24 h prior to transfection. *Control* or *Snrk* siRNA were transfected using Lipofectamine 2000 reagent (Thermo Fisher, Cat# 11668030) for 4-6 h and then changed to complete media and incubated for 48 h until further experimentation such as ELISA or western blot. For co-culturing studies, HL-1 CMs were knockdown for *Snrk* and co-cultured with cardiac FBs. Briefly, HL-1 CMs were grown on the 6-well transwell insert in a separate 6-well plate. *Snrk* (25 μM) and control (25 μM) siRNA was transfected into HL-1 cells with lipofectamine for 4-6 h, followed by a media change to complete media. Then, the transwell inserts were placed on a 6-well plate that contain FBs and incubated for 48 h under co-culture conditions. TGFβR inhibitor (Tocris Bioscience, SB 431542, Cat#1614) at 1 µM concentration or Salmeterol xinofolate (Sigma, Cat#S5068, negative control) at 10 μM was added to cardiac FB wells for 24 h before harvesting the cells for downstream analysis. Cardiac FBs were studied for protein expression by western blot and immunofluorescence.

#### In vitro Immunofluorescence

Immunofluorescence was performed by fixing FBs at 4%PFA (Electron Microscopy Sciences, Cat#15710) for 15 min and washed with PBS and then permeabilized with 0.1% TritonX 100 (BioRad, Cat#1610407) for 15 min and blocked with 4% BSA in PBS for 1 h and overnight incubation of α-SMA antibody (Novus biologicals, Cat#NB300-978) diluted (1:1000) in 1X PBS. Cells were again washed with 1X PBS and incubated with Alexa fluor-488 (Invitrogen, Cat#A11055) diluted (1:500) in 1X PBS for 90 min at RT and washed before mounting with DAPI (LifeSpan Biosciences, Cat#LS-J1033-10) and imaged using a Zeiss confocal microscope at a magnification of ×63. Quantification was done using ImageJ software and plotted as corrected total cell fluorescence (CTCF). CTCF = Integrated Density – (Area of selected cell X Mean fluorescence of background readings).

#### Western blot

Proteins were isolated from HBMECs using Radioimmunoprecipitation Assay buffer (RIPA) buffer (Sigma Cat# R2078) with a complete mini EDTA-free protease inhibitor cocktail (Roche Cat# 11836170001) and PhosSTOP phosphatase inhibitor (Roche Cat# 4906845001). After isolation, the total protein was quantified. Cell lysates were used for probing the following proteins: SNRK (Genetex, Cat# GTX111380, 1:1000 dilution), TGF-β1 (Genetex, Cat# GTX110630, 1:500 dilution), α-smooth muscle actin (Novus Biologicals, Cat# NB300-978, 1:500 dilution), Smad 2/3 (Novus Biologicals, Cat# AF3797, 1:200 dilution), GAPDH (Santa Cruz, Cat# SC-47724, 1:500 dilution) and β-actin (Cell signaling Cat#4970, 1:1000 dilution). The respective primary antibodies were diluted in 1% Tris-buffered saline-Tween20 (TBS-T) and incubated overnight at 4^°^C in the shaker. The primary antibody probed membranes were washed 3X with 1% TBS-T. Secondary HRP antibodies were diluted in 1% TBS-T for 1 h at room temperature in the shaker and washed 3X with 1% TBS-T before adding chemiluminescence detection solution (Pierce™ ECL Western Blotting Substrate, Cat# 32209, ThermoFisher). Anti-rabbit Horseradish peroxidase (HRP) (Cell signaling, Cat#7074, 1:1000 dilution), and anti-goat HRP (Jackson Immunoresearch, Cat#205-052-176, 1:1000 dilution) were secondary antibodies used for chemiluminescence detection. Quantification was done using ImageJ software and plotted against the housekeeping control protein (β-actin) using GraphPad software as described previously (Thirugnanam et al., 2022).

#### ELISA studies

Briefly, HL-1 CM cells were grown in a 30-mm dish and the cells were transfected with *control* or *Snrk* siRNA (25 μM); 100 µL of the cell-free supernatant was used for the assessment of TGF-β1 protein levels. The protocol was followed according to the manufacturer’s instructions (R and D systems Cat# DB100B).

#### Quantification and Statistics

Data were analyzed by two-sample t test, Welch’s t test or analysis of variance (ANOVA) with Tukey’s test to adjust for multiple comparisons. Folded F Test or Levene’s Test were used to examine homogeneity of the variance among different groups. Normality was evaluated by Shapiro-Wilk test. Where necessary for parametric assumptions, log transformation was employed. Mann-Whitney-Wilcoxon test or Kruskal-Wallis test with Dwass, Steel, Critchlow-Fligner Method for multiple comparison adjustment was used where parametric assumptions are not satisfied. P<0.05 was considered significant. SAS version 9.4 (SAS Institute Inc., Cary, NC) was used for statistical analyses.

## Results

### Snrk cardiac conditional knockout mice show more fibrosis in the atrial chamber compared to the ventricular chamber in the basal state

In our previous study, using a non-ischemic Angiotensin II (Ang II)-induced heart failure (HF) mouse model, we showed that SNRK acts as a cardiomyocyte-specific repressor of inflammation and fibrosis (Thirugnanam et al., 2019). When we evaluated fibrosis in a chamber-specific manner in the *Snrk* cardiomyocyte conditional knockout (cmcKO) mice using Sirus red staining for collagen fibers, we observed a slightly higher (**Fig. 1A and A’**, *P<0.05, **P<0.01) fibrosis in the atrial chamber compared to the ventricular chamber in both basal and Ang II infused conditions. Higher magnification section images of the Sirus red stained collagen heart are provided (**Fig. S2A-B**). We also quantified fibrosis around the arteries called perivascular fibrosis and compared them to total tissue between all the groups (**Fig. S2A-C**). We observed significant increase in perivascular fibrosis between the non-Ang II and Ang II treated groups (**Fig. S2C**, P<0.05). This data is clinically relevant in that Ang II levels in atria are increased in patients with Atrial Fibrillation (AF) and HF (Boldt et al., 2003). Ang II also plays a key role in cardiac remodeling and dysfunction in the failing heart (Mehta, and Griendling, 2007; Saito and Berk, 2022). As per the left (LA) and right atrium (RA), we observed variability in the level of fibrosis between the WT (LA: 4.5%; RA: 3.78%), and *Snrk* cmcKO (LA: 25.33%; RA: 21.64%) groups. With Ang II infusion, the WT Ang II (LA: 36.94%; RA: 22.66%) group showed more magnitude of change in LA compared to RA which was not observed in the *Snrk* cmcKO Ang II (LA: 21.81%; RA: 20.51) (**Fig. 1A’’**) group. In addition, we evaluated the expression of alpha smooth muscle actin (α-SMA) in the mouse hearts, where we observed the expression of α-SMA were higher in *Snrk* cmcKO, WT-Ang II and *Snrk* cmcKO Ang II compared to WT (**Fig. S1**). We also investigated the infiltration of pro-fibrotic Mac2 (Galectin3) macrophages (**Fig. 1B and B’**) and did not notice an appreciable difference in their abundance in atria vs. ventricle compartments of *Snrk* cmcKO hearts although there was a total increase in macrophage infiltration into the Ang II-infused and basal *Snrk* cmcKO hearts.

**Fig. 1.**
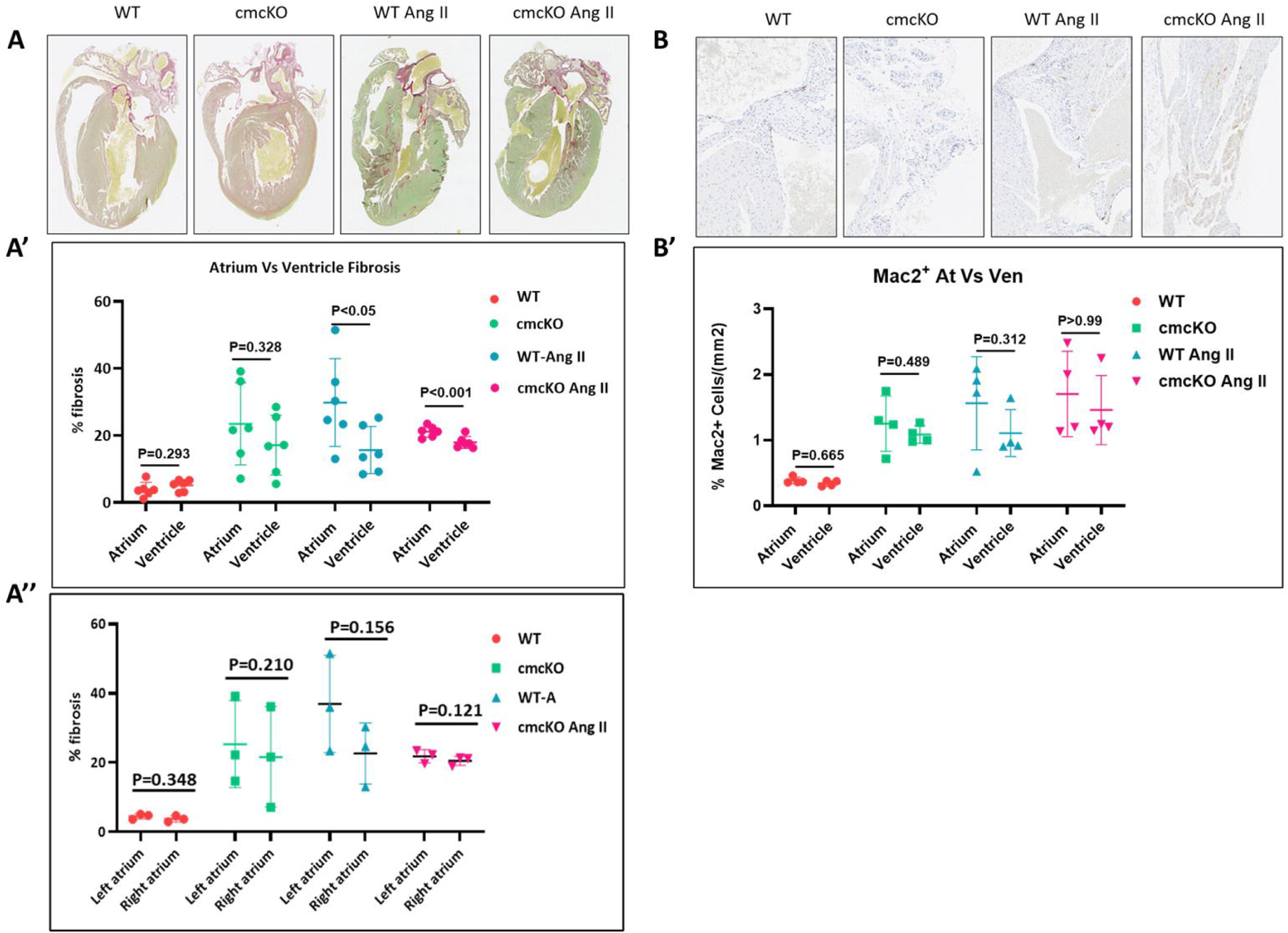
Chamber-specific effect of *Snrk* in cardiac conditional knock-out mice (*Snrk* cmcKO). Experimental conditions include wild-type (WT), cmcKO, WT-Ang II, and cmcKO Ang II. **A and A’.** Sirus red staining of *Snrk* cmcKO hearts was analyzed for chamber-specific levels of fibrosis in the atrium vs. ventricle. Significant increases in the accumulation of collagen in the *Snrk* cmcKO mice atrium were observed in all the experimental conditions. Results are presented as mean ± SD. **A’’.** Quantification of fibrosis in chamber-specific right vs. left atrium. **B and B’.** Mac2^+^ macrophage staining for chamber-specific atrium vs. ventricle macrophage infiltration. Scale bars are 250 μm for *Snrk* WT, *Snrk* cmcKO, and *Snrk* WT-Ang II and 100 μm for *Snrk* cmcKO-Ang II. Results are presented as mean ± SD. The analysis and statistical representation of the data were only from male mice, n=3 for WT and n=3 for conditional knockout groups. The 6 data points represented per condition in the bottom panel were one from the left and right atrium and one from the left and right ventricle per mice (n=3). The ages of the mice for the wild type and *Snrk* cmcKO group range from 3 months, 5 days (youngest) to 4 months, 18 days (oldest). All data sets were normalized with body weight measurements. For panels 1A-A’, ANOVA statistical tests was done. For panels 1B-B’, Wilcoxon statistical test was done.

### Higher Snrk expression in atria is conserved in rats and humans and is associated with preventing atrial fibrosis and its related effects in HF

To evaluate whether SNRK showed selective expression in atrial vs. ventricular cells in the heart, we performed IF for SNRK in rat hearts (**Fig. 2A-B**). Confocal IF images for SNRK protein in atria and ventricle of a 56-day old rat heart showed more SNRK expression in LA than in LV (LA: 1.1±0.4 vs. LV: 0.5±0.1 [cells/µm²], P=0.055, n=3) (**Fig. 2C**). To determine SNRK expression in cardiomyocytes (CMs) in both chambers, we co-stained for cardiac troponin T (TnT) along with SNRK in LA and LV (**Fig. 2D-F**). Indeed, in control rats, SNRK-positive CMs in LA chamber was 21.4 ± 3.9% compared to 10.1 ± 1.4% in LV chamber (**Fig. 2F**). Upon Ang II infusion, SNRK expression in LA CMs went down to 6.3 ± 4.2% from 21.4% in controls, and in LV CMs, went down from 10.1% to 6.3% ± 3.1% (**Fig. 2F**). Thus, more atrial CMs express SNRK, and upon Ang II infusion, the magnitude of change in SNRK expression was greater in LA vs. LV CMs.

**Fig. 2.**
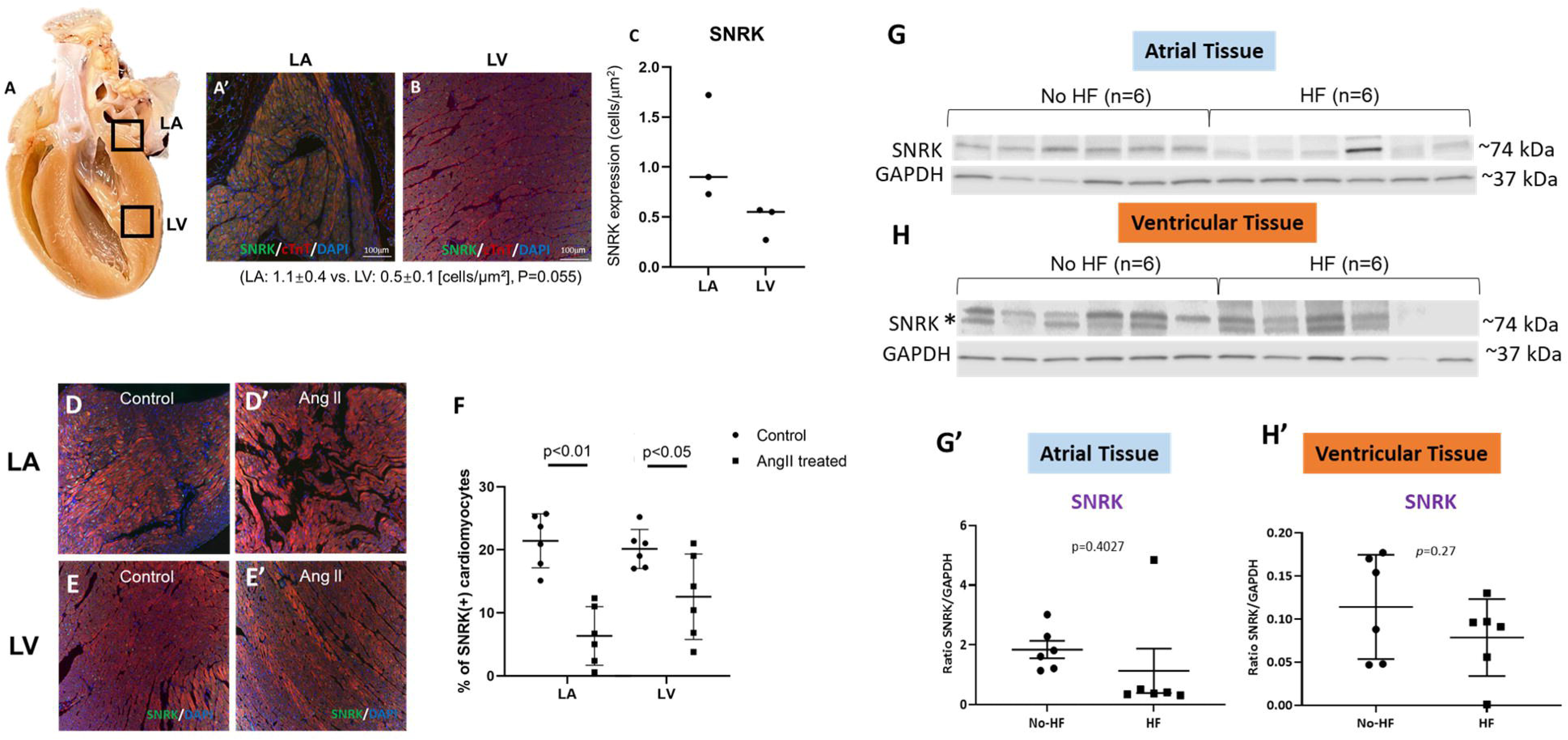
Chamber and septum-specific expression of SNRK levels. **A-B. A** shows the location of the rat heart where the images were taken for left atria (LA) and left ventricle (LV). **A’-B** are merged (SNRK, cardiac troponin (cTnT) and DAPI) immunofluorescence images of the septum-specific LA and LV. **C.** Quantification of SNRK levels in the septum-specific LA vs. LV. Results are presented as mean ± SD. N=3. **D-F.** Immunofluorescence analysis of TnT co-stained with SNRK in septum-specific LA vs. LV cardiomyocytes treated with and without Angiotensin II (Ang II). Results were quantified and presented as mean ± SD. N=3. **G-H’.** Atrial and ventricular human heart failure and non-heart failure patient samples were analyzed for SNRK protein levels by immunoblotting method. The bottom band for SNRK protein marked with an asterisk (*) was quantified. Results were quantified and presented as mean ± SD. N=6 per condition. Control blots for the housekeeping protein GAPDH were stripped and re-probed for SNRK. For all panels, ANOVA test was done.

To assess the relevance of SNRK expression in human tissue, heart tissue from six patients each (HF and no HF) undergoing procedures (ventricular assist device implantation or cardiac transplant) at Aurora, St. Luke Medical Center (Milwaukee, WI) were collected as part of an existing IRB at Advocate Aurora Health, Milwaukee, WI (**Fig. S3G**). The HF samples were verified for increased CD68 macrophage staining and increased collagen staining for fibrosis (**Fig. S3**) compared to no HF samples. Atrial (**Fig. 2G, G’**) and ventricular (**Fig. 2H, H’**) lysates from patients with HF and no HF were generated and probed for SNRK and GAPDH proteins. We observed that SNRK expression in atrial tissue was higher compared to ventricular tissue in no HF patient (notice *y-axis* for no HF samples, **Fig. 2G’-H’**). Note that the SNRK antibody detected two bands in ventricular heart tissue compared to atrial tissue. Both bands were quantified. The lower band (black asterisk) that matched SNRK size (~74 kDa) and the upper band (**Fig. S4A**), showed appreciable differences across samples (HF and no HF groups). Also, SNRK expression is lost in 5 out of 6 HF atria samples (**Fig. 2G**). Taken together, data in rats and humans suggest that SNRK expression is markedly higher in atrial CMs compared to ventricle CMs and may have functional implications in preventing AF or HF associated with AF.

### SNRK in atrial cardiomyocyte mediates repression of transforming growth factor-β1 expression

To investigate the underlying mechanism of how SNRK in CMs prevents fibrosis, we focused on transforming growth factor-β1 (TGF-β1) as a potential target of SNRK because TGF-β1 was previously implicated in promoting selective atrial fibrosis in a transgenic overexpressing mouse system (Nakajima et al., 2000). We investigated whole heart tissue lysates for TGF-β1 protein levels in the vehicle and Ang II infused *Snrk* cmcKO mice (**Fig. 3A**) and in control rat hearts with and without Ang II (**Fig. 3C**). TGF-β1 protein levels were higher in basal *Snrk* cmcKO heart (**Fig. 3B**) and upon Ang II, these levels continued to rise even further (**Fig. 3B**). In rats with Ang II infusion for 28 days, TGF-β1 was immunostained in rat atria and ventricle CMs and were quantified. TGF-β1 expression is higher in Ang-II-infused LA CMs compared to Ang-II-infused LV CMs (**Fig. 3D**). In human atrial tissue, TGF-β1 protein levels also showed a similar trend towards higher expression in HF over no HF patient samples (**Fig. 3E**). Because TGF-β1 protein is secreted, we probed for TGF-β1 protein via ELISA in control and efficacy-confirmed *Snrk* siRNA knockdown HL-1 adult immortalized mouse atrial cell supernatants (**Fig. 3F**). We detected 48 pg/mL of TGF-β1 in *Snrk* siRNA compared to control siRNA of 7 pg/mL. The enhanced TGF-β1 protein levels detected by ELISA was also confirmed by western blots of *Snrk* siRNA HL-1 cell lysates (**Fig. 3G**). These expression studies collectively suggest that SNRK regulates TGF-β1 expression in atrial CMs.

**Fig. 3.**
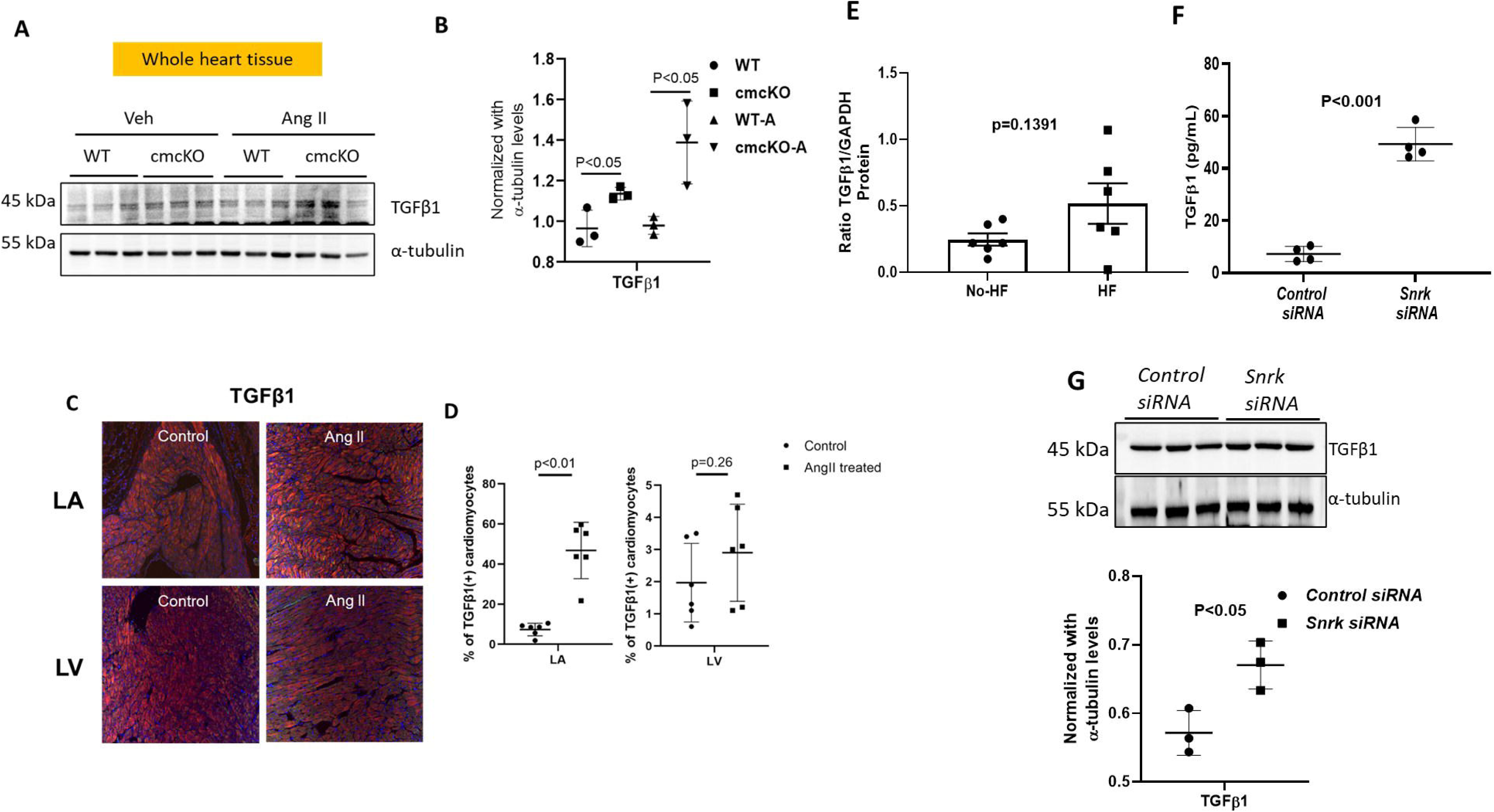
Cardiomyocyte SNRK represses TGFβ1 signaling. **A.** TGFβ1 protein immunoblotting in whole heart tissue lysates from wild type (WT) and *Snrk* cmcKO mice with and without Angiotensin II (Ang II) treatment. **B.** Quantification of TGFβ1 protein expression from the blot in **A**. Results are presented as mean ± SD. N=3. Control blots for the housekeeping protein α-tubulin were stripped and re-probed for TGFβ1.**C-D.** Immunofluorescence analysis of the expression of TGFβ1 in the septum-specific left atrium (LA) vs. left ventricle (LV) cardiomyocytes with and without Ang II. Results are presented as mean ± SD. N=3. **E.** Human heart failure and non-heart failure atrial patient samples were analyzed for TGFβ1 protein levels by immunoblotting method. Results are presented as mean ± SD. N=6. **F.** Mouse atrial HL-1 CMs knockdown for *Snrk* and assessed for TGFβ1 protein levels in the cell-free supernatant method by ELISA. Results are presented as mean ± SD. N=3. **G.** Mouse atrial HL-1 CMs knockdown for *Snrk* and assessed for TGFβ1 protein levels by immunoblotting method. Results are presented as mean ± SD. N=3. Control blots for the housekeeping protein α-tubulin were stripped and re-probed for TGFβ1. For panels 3B, 3F & 3G t-test was done. For panels 3D & 3E, Welch’s t-test was done.

### TGF-β1 released from SNRK KD atrial CMs promotes fibrosis

TGF-β1 is a well-studied molecule that plays a critical role in cardiac fibrosis events (Nakajima et al., 2000; Li et al., 2008; Tsai et al., 2011; Zeisberg et al., 2007). To investigate the direct role of TGF-β1 secreted from *Snrk* knockdown atrial CMs for cardiac fibrosis, we performed co-culture studies using transwell inserts. HL-1 atrial CMs knocked down for *Snrk* co-cultured with mouse cardiac fibroblasts (FBs) were studied for FBs activation with and without TGFβR inhibitor or negative control Salmeterol xinafoate in FBs. The protein expression studies suggest that the knockdown of *Snrk* in CMs increases the expression of α-smooth muscle actin (α-SMA) and the Smad2/3 pathway in FBs. Whereas the *Snrk* knockdown CMs co-cultured with fibroblasts treated for TGFβR inhibitor showed a significant decrease in the expression of α-SMA and the Smad2/3 proteins (**Fig. 4A-D**). The results from HL-1 atrial CM-mediated *Snrk* knockdown co-cultured FBs by immunofluorescence also suggest a significant increase in the activation of α-SMA in FBs (**Fig. 4E-F**). Collectively, the co-culture experiment data suggests that SNRK associated TGFβ1 secreted from CMs plays a significant role in the activation of FBs, leading to myofibroblast and fibrotic phenotype.

**Fig. 4.**
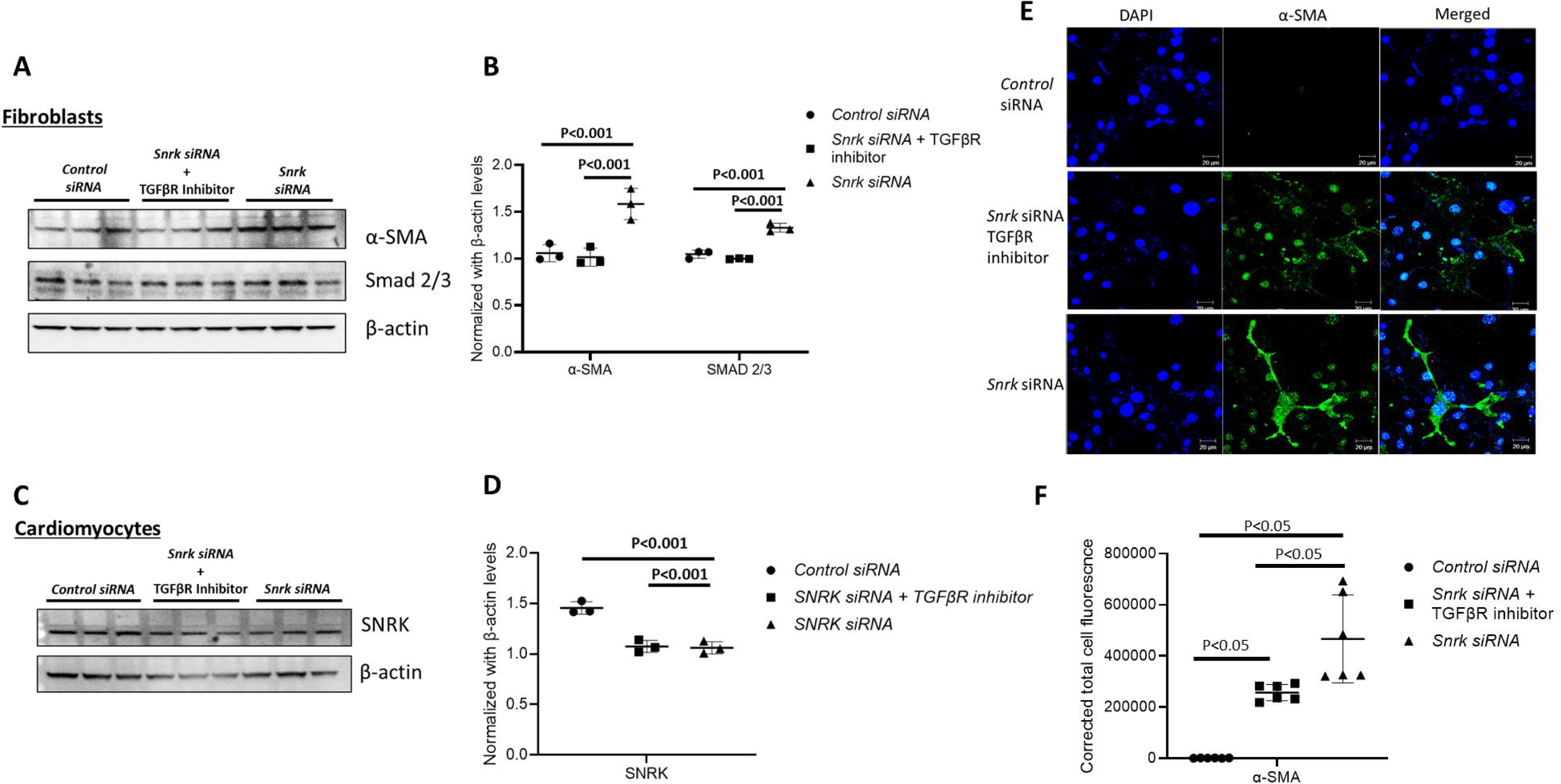
SNRK cardiomyocyte repressed TGFβ1 prevents fibroblast activation. **A-B.** Mouse atrial HL-1 CMs knockdown for *Snrk* and co-cultured with cardiac fibroblasts were assessed for the expression of fibroblast activation markers, Smad 2/3 and α-SMA by immunoblotting method and quantified respectively. The experiment was performed with and without TGFβR inhibitor (1 µM) or negative control Salmeterol xinafoate on the cardiac fibroblasts. Results are presented as mean ± SD. N=3. Control blots for the housekeeping protein, β-actin were stripped and re-probed for α-SMA and SMAD 2/3. **C-D.** Mouse atrial HL-1 CMs knockdown for *Snrk* and assessed for knockdown efficiency. Results are presented as mean ± SD. N=3. **E-F.** Mouse atrial HL-1 CMs knockdown for *Snrk* and co-cultured with cardiac fibroblasts were assessed for the expression of α-SMA by immunofluorescence. Quantification was done in ImageJ to calculate the corrected total cell fluorescence (CTCF). CTCF = Integrated Density – (Area of selected cell X Mean fluorescence of background readings). Quantification was done on two random areas per slide per condition. Results are presented as mean ± SD. N=3. Scale bar= 20 µm. Control blots for the housekeeping protein β-actin were stripped and re-probed for SNRK. For panels 4B & 4D, ANOVA with Tukey adjustment test was done. For panel 4F, Kruskal-Wallis test with Dwass, Steel, Critchlow-Fligner Method was used.

## Discussion

In this study, we identified SNRK in atrial CMs as a potential cardiac repressor of TGFβ1 signaling, an important regulator of cardiac fibrosis. Previously, we demonstrated cardiac functional deficits, inflammation, and fibrosis as major phenotypes in the heart of *Snrk* cmcKO mice (Thirugnanam et al., 2019; Cossette et al., 2016; Cossette et al., 2014). In the current study, the chamber-specific analysis of fibrosis revealed (**Fig. 1 and S1**), that *Snrk* cmcKO hearts exhibit a higher level of fibrosis in the atria compared to the ventricle (**Fig. 1A**). The chamber-specific trend observed in the *Snrk* cmcKO was also comparable to the Ang II-infused mice suggesting the preferential occurrence of fibrosis in the atria. Within the atrium, the WT did not express high variability as they have minimal fibrosis, and in *Snrk* cmcKO and WT Ang II condition, there is a difference between left atrium (LA) and right atrium (RA), although they are not statistically significant. In the *Snrk* cmcKO Ang II condition, the fibrosis is evenly distributed between the LA and RA. The important point here is, that these *Snrk* cmcKO Ang II mice die within 14 days of Ang II infusion, which implies that these *Snrk* cmcKO Ang II mice are on the verge of dying, and their cardiac system is compromised, and the amount of fibrotic deposition are perhaps reaching a saturation point. Further, it is intriguing to note that when the fibrosis in the LA is higher, although the cardiac functional parameters are declining, the mice did not die immediately (*Snrk* cmcKO mice), but when the fibrosis in the RA is increased and equivalent to the LA, they die rapidly (*Snrk* cmcKO Ang II mice). Therefore, this raises the question of crosstalk compensatory mechanism between the LA and RA, which we will be investigated in future studies. Our current cardiac fibrosis data (~3 months) when compared to previous studies (~6 months), showed a 2.4-fold increase suggesting an age-dependent progression of fibrosis in the *Snrk* cmcKO mice.

In this study, we deleted *Snrk* from all CMs. Thus, we cannot eliminate the contribution of ventricular CMs to fibrosis. Our previous publication (Thirugnanam et al., 2019) showed ejection velocity changes in ECHO in *Snrk* cmcKO mice. Defective ejection velocities of blood are common during cardiac relaxation and contractility (Thirugnanam et al., 2019) a hallmark event in ventricular fibrosis (Alhakak et al., 2021). Thus, we cannot exclude an indirect effect of *Snrk* loss on ventricles as a factor for fibrosis progression and cause of HF. An atrial-specific CRE excision of SNRK is necessary to bolster our conclusions and the lack of this approach is a limitation of our study. In terms of macrophage infiltration, the Mac2^+^ macrophage infiltration does not show any chamber-specific effect other than the basal level increase in *Snrk* cmcKO and Ang II treated mice compared to the WT condition (**Fig. 1B**). This could imply that in the failing heart, the persistent migration of chamber-specific Mac2^+^ cells may be saturating as the mice are already proceeding toward progressive fibrosis and on the brink of death.

IF analysis of septum-specific SNRK reveals that SNRK is one-fold higher in the left atrium compared to the left ventricle under normal baseline conditions (**Fig. 2A-C**). In terms of CMs, IF analysis of SNRK in septum-specific CMs suggests, that SNRK is one-fold higher in the left atrial CMs than in the left ventricular CMs (**Fig. 2D-F**). The preferential atrial SNRK protein expression is also recapitulated in human hearts. SNRK protein analysis on human heart samples for HF and non-HF reveals that SNRK is significantly higher in the atrium than the ventricle in the non-HF samples (**Fig. 2G-H**). In addition, compared to non-HF samples, SNRK expression is decreased in HF patients. All these results collectively suggest the hypothesis that SNRK expression in left atria is responsible for repressing fibrosis in atria. A recent single-cell analysis of the human heart tissue reveals, composition-wise, the atrium contains 30.1% of CMs and 24.3% of FBs compared to 49.2% of CMs and 15.5% of FBs in the ventricle (Litviňuková et al., 2020). Taken together, the presence of more FBs in atria may increase the propensity for fibrosis in the atrium, and the significant increase in the expression of SNRK in the atrium may provide important regulation of atrial FBs in the normal homeostatic functioning of the heart. Our previous study also showed that the endothelial-specific *Snrk* knockout mice had inflammation but did not progress to fibrosis, which suggests compensation mechanisms promoted by CM SNRK (Thirugnanam et al., 2019). The current data supports this hypothesis and suggests increased expression of SNRK in the atrial CMs as a protective checkpoint in the heart. In this hypothesis, in HF, when SNRK levels plummet in the atrium (**Fig. 2 G-H**), the protective checkpoint is lost leading to enhanced fibrosis and a failing heart. However, how the left atria and right atria CMs communicate with chamber-specific FBs and with other cells in the heart, and SNRK’s role in that is not known and will need further investigation.

In terms of SNRK regulation in fibrosis signaling, we assessed TGFβ1 levels. TGFβ1 is a prominent regulator of inflammatory, hypertrophic, and fibrotic processes in myocardial remodeling caused by various cardiac stress and disease forms (Rainer et al., 2014). The data from mice, rats, human hearts, and in vitro mouse atrial CMs reveals, that a loss or decrease in CM SNRK levels, increases the expression of TGFβ1 levels (**Fig. 3**) suggesting SNRK acts as a checkpoint in regulating TGFβ1. The septum-specific data from the rats suggests the TGFβ1 levels are higher in the atrium compared to the ventricle with Ang II treatment or even baseline levels (**Fig. 3C and D**). The atrium-specific increase in the baseline levels of SNRK or TGFβ1 is an important finding of this study. TGF-β1 activates fibroblasts and stimulates them to differentiate into myofibroblasts, which produce extracellular matrix (ECM) proteins and induce fibroblast proliferation (Saadat et al., 2020). A previous study suggests that cardiomyocyte-driven mutant TGFβ1 expressing mice show only atrial fibrosis although the constitutive active protein is found in both chambers (Ponikowski et al., 2014). This data along with ours suggests that the atrium is highly sensitive to any cardiac stress or inflammatory insults. In support of the atrial-specific fibrosis, the Smad 2/3 and α-SMA expression in cardiac fibroblasts from the co-culture studies with western blot and immunofluorescence confirm that SNRK-mediated TGFβ1 secreted from CMs is sufficient to induce fibroblast activation to myofibroblast (**Fig. 4**). SMA expression was also upregulated in *Snrk* cmcKO mice and these levels went even higher upon Ang II infusion (Fig. S1). The activation of cardiac fibroblast to myofibroblast is the key process in the event of cardiac fibrosis. The myofibroblast activation is commonly associated with most cardiovascular disease (Liu et al., 2021). The activation of the Smad pathway in myofibroblasts, recruits and activates intracellular effectors to send the signal to the nucleus, which triggers gene transcription for ECM components such as α-SMA towards fibrosis (Vallee et al., 2019). Targeted disruption of TGFβ receptor signaling improves survival after myocardial infarction (Rainer et al., 2014). Indeed, TGFβ receptor inhibition in cardiac FBs co-cultured with *Snrk* KD CMs revealed that secreted TGFβ1 from CMs were unable to signal to FBs, which was assessed by a decrease in the expression of Smad 2/3 and α-SMA proteins in FBs. Thus, we conclude that SNRK associated TGFβ1 atrial CMs-FB signaling leads to myofibroblast and fibrotic phenotype.

In summary, we have identified an atrial selective target SNRK that is expressed more in atria over the ventricle in multiple species and helps prevent the secretion of a potent myofibroblast driver TGFβ1 to prevent atrial fibrosis selectively in atrial CMs. Such a chamber-specific SNRK-TGFβ1 axis may be highly beneficial for developing targeted approaches to prevent atrial fibrosis in non-ischemic HF.

## Data availability statement

The raw data supporting the conclusion of this article will be made available by the authors, without undue reservation.

## Ethics statement

The studies involving animals followed Animal Care and Use Committee Guidelines and were approved by the respective institutions accordingly as mentioned in the methods section.

## Supporting information

All supplemental figures

## Acknowledgements

We thank the Department of Pediatrics at the Medical College of Wisconsin, Children’s Research Institute at Children’s Wisconsin for supporting the Developmental Vascular Biology program with programmatic funds that partly facilitated this research. RR, KT and SP are partly supported by R61HL154254. RR and KT are also supported by program development funds from Children’s Research Institute. FR, AJ and PH were partly supported by RO1 HL101240 from NHLBI/NIH. FR was also supported by 570-5038, Aurora Cardiovascular Surgery Research Award. XB was supported by grants R01 GM112696 and 1R35GM148177 from the NIH, Advancing a Healthier Wisconsin, and Medical College of Wisconsin-Neuroscience Research Center Award. HS was supported by JSPS KAKENHI Grant Number JP21K20415. We thank the National Disease Research Interchange (NDRI) for the human heart samples.

## Publisher’s note

All claims expressed in this article are solely those of the authors and do not necessarily represent those of their affiliated organizations, or those of the publisher, the editors, and the reviewers. Any product that may be evaluated in this article, or claim that may be made by its manufacturer, is not guaranteed, or endorsed by the publisher.

## Supplementary Figure Legends

**Fig. S1. Mouse heart tissue section assessed for the expression of alpha smooth muscle actin.** Wild type, *Snrk* cmcKO, Wild type-Ang II and *Snrk* cmcKO-Ang II tissue sections stained for α-SMA and quantified. Results are presented as mean ± SD. N=3. Statistical analysis (P value) was compared to WT. Mann-Whitney-Wilcoxon statistical test was done.

**Fig. S2. Representative images of whole heart section for Sirius Red fibrosis staining. A**. Wild type and *Snrk* cmcKO whole heart section. **B.** Wild type-Ang II and *Snrk* cmcKO-Ang II whole heart section. **C.** Whole heart tissue samples were quantified for perivascular fibrosis and compared to the total tissue area. Results were presented as mean ± SD. N=3. Mann-Whitney-Wilcoxon statistical test was done.

**Fig. S3. Human heart failure and non-heart failure samples assessed for inflammation and fibrosis. A-C.** HF and no-HF samples were stained for collagen to assess fibrosis and quantified with the ratio of collagen-positive area to myocardial area. **D-F.** HF and no-HF samples were stained for CD68-positive macrophages and quantified per mm^2^. **G**. Representative table of human patient sampling data. For panels, C & F, t-test was done.

**Fig. S4. A.** Human heart failure and non-heart failure ventricular samples were quantified for the top band in figure 2H. **B.** Salmeterol xinafoate (10 μM) treated mouse cardiac fibroblasts were assessed for α-SMA. Salmeterol xinafoate is a negative control. Scale bar: 20 μm. For panel A, t-test was done.

