## Supplementary material for "SNRK regulates TGFβ levels in atria to control cardiac fibrosis": All supplemental figures

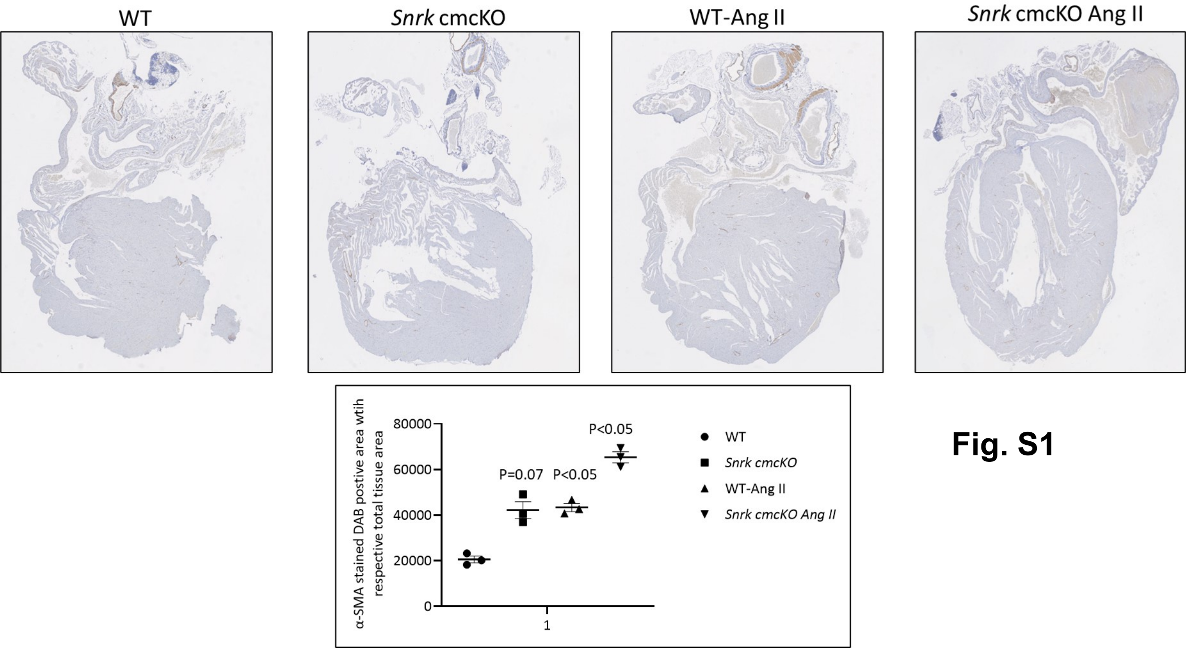
WT
Snrk cmcKO
WT-Ang II
Snrk cmcKO Ang II
α-SMA stained DAB postive area wtih respective total tissue area
80000
60000
40000
20000
0
1
P=0.07
P<0.05
P<0.05
WT
Snrk cmcKO
WT-Ang II
Snrk cmcKO Ang II

**Fig. S1**

**Fig. S2A**

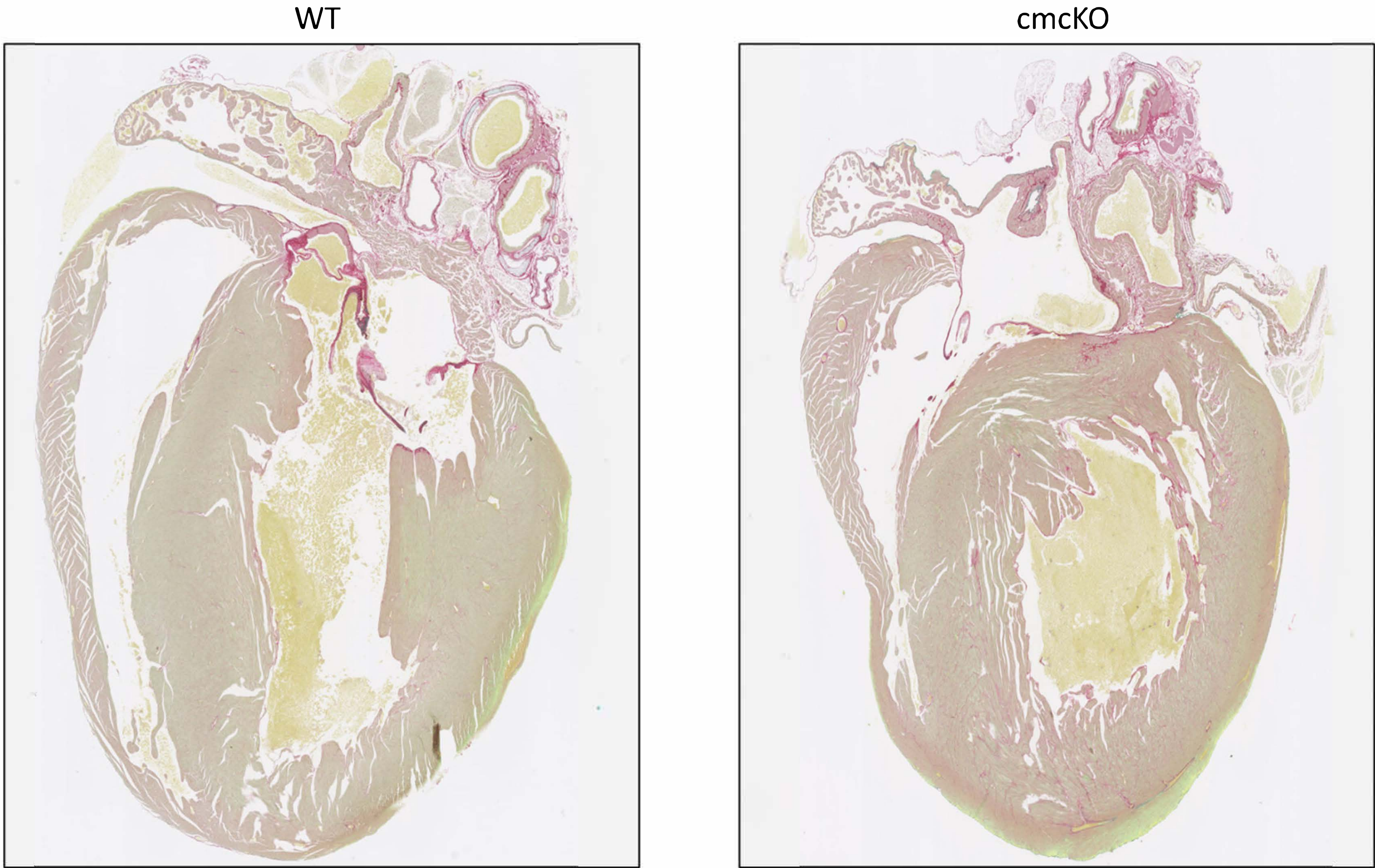

**Fig. S2B**

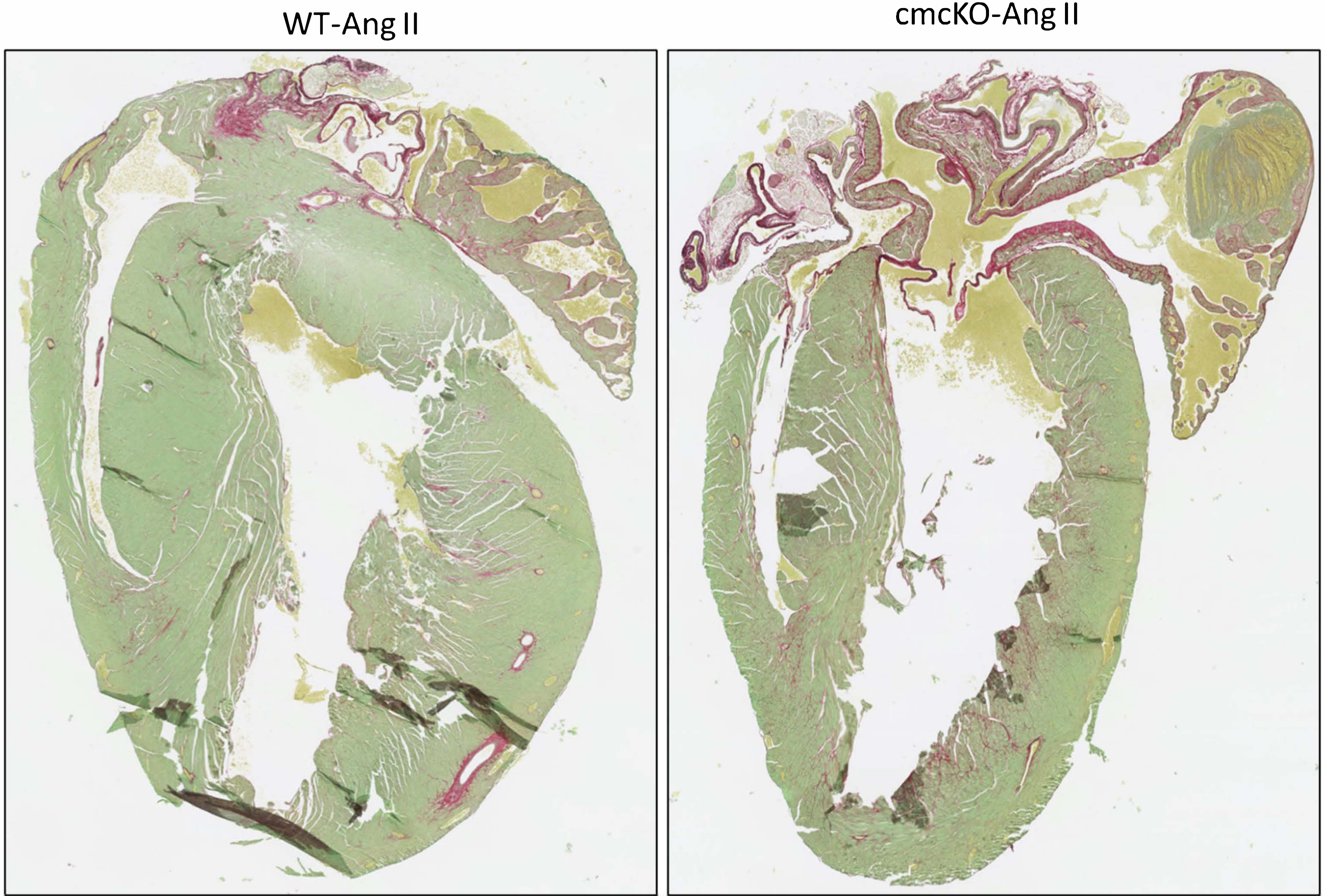


**Fig. S2C**

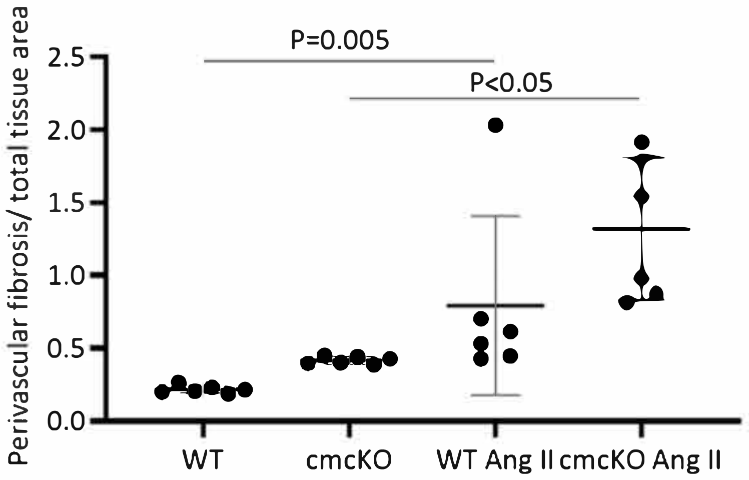

**Fig. S3**

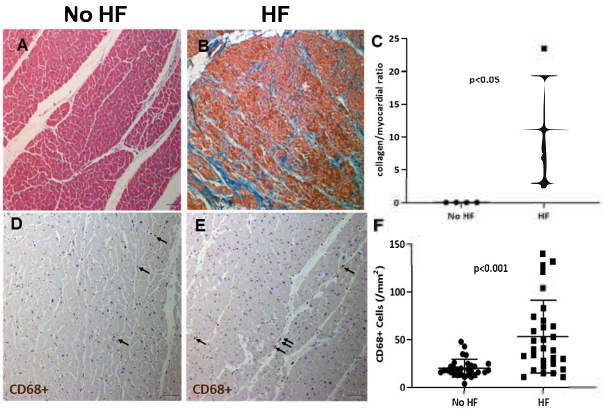


**G**

| Sample ID | Sample type | Disease/underlying conditions | Age (Y) | Gender | HFpEF/HFrEF |
|---|---|---|---|---|---|
| **Heart Failure Samples** | | | | | |
| ND-1 | Donor Dead | HF/Diabetes/Afib | 81 | M | N/A |
| HT23 | Heart Transplant | HF/Cancer | 62 | M | HFrEF |
| HT24 | Heart Transplant | HF/Diabetes | 67 | M | HFrEF |
| HT25 | Heart Transplant | HF/Diabetes | 32 | F | HFrEF |
| HT26 | Heart Transplant | HF/Diabetes | 42 | M | HFrEF |
| HT29 | Heart Transplant | HF/Cancer | 64 | M | HFrEF |
| **Non-Heart Failure Samples** | | | | | |
| ND-2 | Donor Dead | Hyperlipidemia/COPD | 78 | F | N/A |
| ND-3 | Donor Dead | Hyperlipidemia/GERD/Neuropathy | 50 | F | N/A |
| ND-5 | Donor Dead | Renal Failure | 45 | M | N/A |
| ND-7 | Donor Dead | Diabetes/Hyperlipidemia | 80 | F | N/A |
| ND-8 | Donor Dead | Hyperlipidemia/GERD | 64 | M | N/A |
| ND-9 | Donor Dead | Alzheimer | 81 | F | N/A |

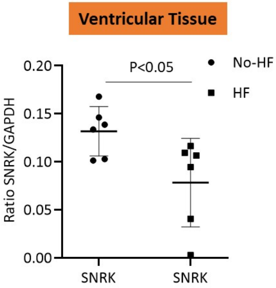

Ventricular Tissue
0.20
0.15
0.10
0.05
0.00
Ratio SNRK/GAPDH
P<0.05
No-HF
HF
SNRK
SNRK


**Fig. S4**

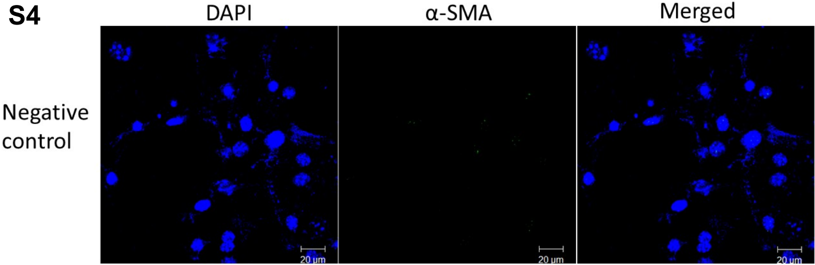

DAPI
α-SMA
Merged
Negative control
20 µm
20 µm
20 µm
